# The *Pax6* master control gene initiates spontaneous retinal development via a self-organising Turing network

**DOI:** 10.1101/583807

**Authors:** Timothy Grocott, Estefania Lozano-Velasco, Gi Fay Mok, Andrea E Münsterberg

## Abstract

Understanding how complex organ systems are assembled from simple embryonic tissues is a major challenge. Across the animal kingdom a great diversity of visual organs are initiated by a ‘master control gene’ called *Pax6*, which is both necessary and sufficient for eye development^1–6^. Yet precisely how *Pax6* achieves this deeply homologous function is poorly understood. Here we show that vertebrate *Pax6* interacts with a pair of morphogen-coding genes, *Tgfb2* and *Fst*, to form a putative Turing network^7^, which we have computationally modelled. Computer simulations suggest that this gene network is sufficient to spontaneously polarise the developing retina, establishing the eye’s first organisational axis and prefiguring its further development. Our findings reveal how retinal self-organisation may be initiated independent of the highly ordered tissue interactions that help to assemble the eye *in vivo*. These results help to explain how stem cell aggregates spontaneously self-organise into functional eye-cups *in vitro*^8^. We anticipate these findings will help to underpin retinal organoid technology, which holds much promise as a platform for disease modelling, drug development and regenerative therapies.

## Introduction

Positional cues that govern cell fate decisions in the embryo may arise at multiple organisational levels: cell-intrinsically (e.g. asymmetric cell divisions), tissue-intrinsically (e.g. reaction-diffusion mechanisms), tissue-extrinsically (e.g. inductive tissue interactions) or some combination of these. Historically, the early patterning of cell fates within the vertebrate eye has emphasised inductive interactions, stemming from Spemann’s seminal work on lens induction^9^. These inductive interactions coordinate self-assembly of the various tissues that comprise the vertebrate camera eye including the optic vesicle of the forebrain, which generates the retina, and the overlying presumptive lens tissue. In the embryo, interactions with neighboring tissues help to remodel the hemi-spherical optic vesicle into a bi-layered optic cup. Yet this vesicle-to-cup transformation is spontaneously recapitulated by stem cell-derived retinal organoids in vitro^8^, revealing that a hitherto unsuspected tissue-intrinsic mechanism suffices to self-organise the primary retinal axis. Here we provide evidence for a self-organising mechanism centered on the transcription factor-coding gene *Paired box 6* (*Pax6*).

*Pax6* has been called an eye master control gene^10^ and is necessary for eye development across much of the animal kingdom, from flies to humans^1–3, 11^. Mis-expression of mammalian or cephalopod *Pax6* genes triggers the spontaneous development of ectopic compound eyes in arthropods^4, 5^, as well as supernumerary camera eyes in vertebrates^6^. This deeply homologous function, whereby a shared *Pax6* genetic apparatus builds eye structures that are morphologically and phylogenetically distinct^12^, is poorly understood. Here we describe a putative self-organising Turing network^7^ comprising *Pax6* and a pair of morphogen-coding genes *Transforming Growth Factor-beta 2* (*Tgfb2*) and *Follistatin* (*Fst*). Using reaction-diffusion modelling we show how this gene network may spontaneously polarise the optic vesicle to trigger self-organisation of the vertebrate retina.

## Results

Optic vesicle polarisation is apparent from Hamburger & Hamilton^13^ stage HH10 in the chick, evidenced by differential gene expression along a proximal-distal axis: *Pax6* and *Visual system homeobox 2* (*Vsx2*; formerly *Chx10*) are expressed distally (Fig.1a, b), whereas *Microphthalmia associated transcription factor* (*Mitf*) and *Wnt family member 2b* (*Wnt2b*; formerly *Wnt13*) are expressed proximally (Fig.1c, d). We additionally report that two further genes, *Transforming Growth Factor-beta 2* (*Tgfb2*) and *Follistatin* (*Fst*) are co-expressed with *Pax6* in the distal optic vesicle (Fig.1e, f).

As the optic vesicles evaginate between stages HH8 and HH10, they encounter Bone morphogenetic protein (Bmp) family growth factors from the overlying surface ectoderm (e.g. *Bmp4*; Fig. 1g). Bmps are implicated in establishing both distal and proximal cell identities within the optic vesicle; Bmp alone promotes distal character^14^, whereas combined with canonical Wnt signalling it was proposed to induce proximal character^15^. Consistently, we found that exposing HH10 optic vesicle explants to Bmp4 ligand for 16 hours *in vitro* led to an up-regulation of distal *Pax6* (2.35 ± 0.19 fold, mean ± standard deviation; *P* < 0.01; *n* = 4) as measured by RT-QPCR (Fig. 1h). The remaining distal (*Vsx2*) and proximal (*Wnt2b, Mitf*) markers were not significantly affected (Fig. 1h). Following combined exposure to both Bmp4 and the Wnt agonist BIO (6-bromoindirubin-3’-oxime; GSK3 inhibitor)^16^, *Pax6* (1.88 ± 0.38 fold; *P* < 0.05; *n* = 5) was similarly affected (Fig. 1i), while the proximal marker *Wnt2b* was additionally up-regulated (9.28 ± 7.89 fold; *P* < 0.05; *n* = 5), suggesting that *Wnt2b* may auto-regulate. Wnt activation alone induced proximal *Wnt2b* (3.69 ± 1.43 fold; *P* < 0.01; *n* = 4) without significantly affecting distal markers (Fig. 1j), while exposure to DMSO (carrier for BIO) had no impact (Fig. 1k). These data do not support a direct synergism between Bmp and Wnt signalling in establishing proximal-distal polarity, as their combined action is merely additive.

**Figure 1:**
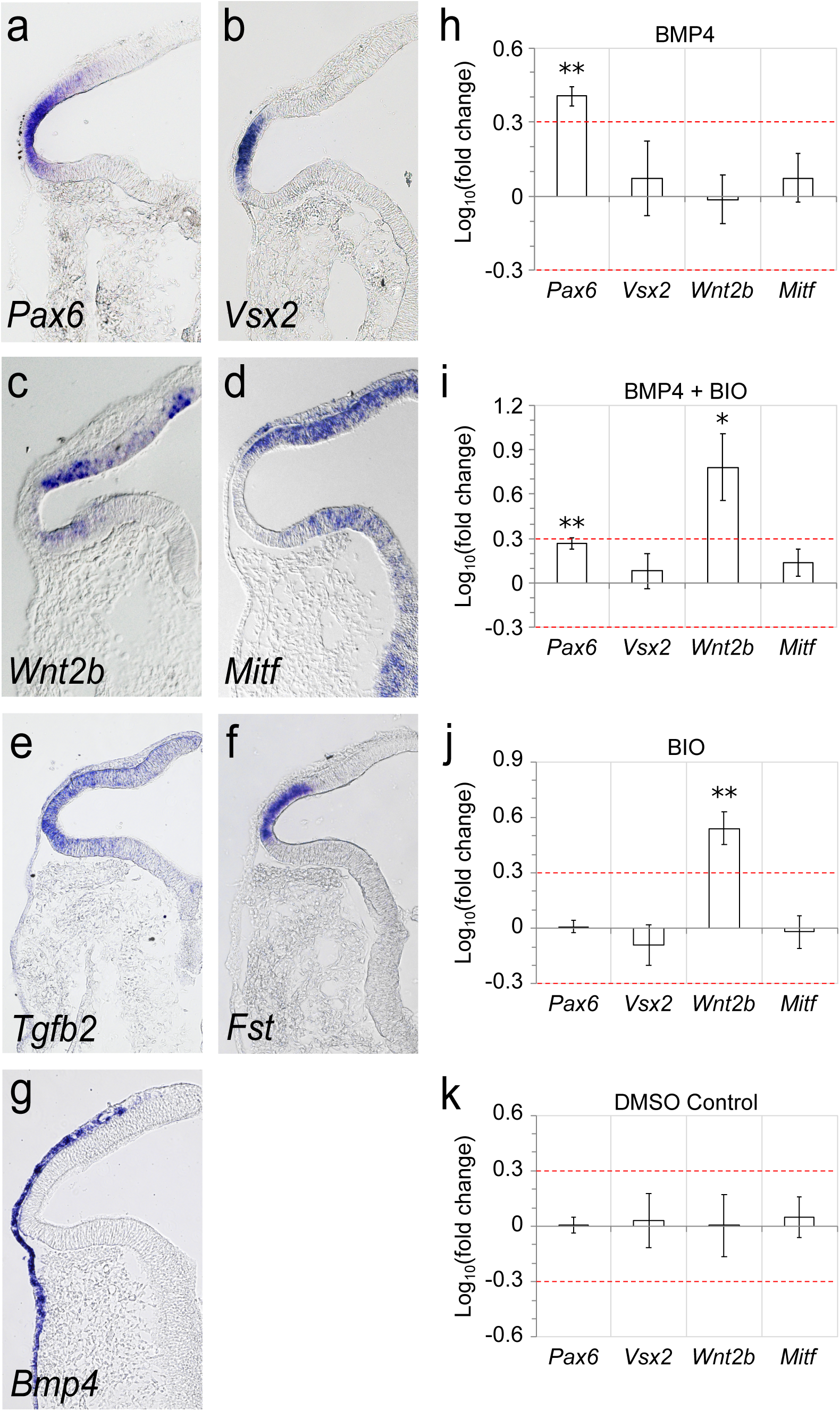
Bmp and canonical Wnt signalling do not directly synergise to induce proximal identity in the optic vesicle. **a-f)** The HH10 optic vesicle is polarised along a proximal-distal axis. Horizontal sectioning reveals polarised expression of the marker genes **a)** *Pax6*; **b)** *Vsx2*; **c)** *Wnt2b*; **d)** *Mitf*; **e)** *Tgfb2*; **f)** *Fst*. **g)** *Bmp4* is expressed in the overlying presumptive lens ectoderm. **h-j)** RT-QPCR analysis of proximal and distal marker gene expression following 16-hour exposure to **h)** Bmp4 only; **i)** Bmp4 and BIO (a canonical Wnt agonist) in combination; **j)** BIO only; **k)** DMSO carrier control. Values plotted are Log10(mean fold change) +/-SEM. Red guide lines indicate the levels of +/-2-fold change in gene expression. * P < 0.05; ** P < 0.01.

To validate the interaction between Bmp signalling and *Pax6* expression *in vivo*, we performed electroporation-mediated gene transfer to mis-express the cell-autonomous Bmp inhibitor *Smad6* in single optic vesicles, while un-electroporated contralateral vesicles served as internal negative controls (Fig. 2a). In comparison to mis-expression of a benign Enhanced Green Fluorescent Protein (GFP; 1.13 ± 0.37 fold; *n* = 7; Fig. 2b, c, d), *Smad6* caused a asymmetric reduction in the area of *Pax6* expression between transfected and contralateral control vesicles (0.56 ± 0.31 fold; *P* < 0.05; *n* = 13; Fig. 2b, e, f). This confirms that distal *Pax6* expression *in vivo* requires upstream Bmp.

**Figure 2:**
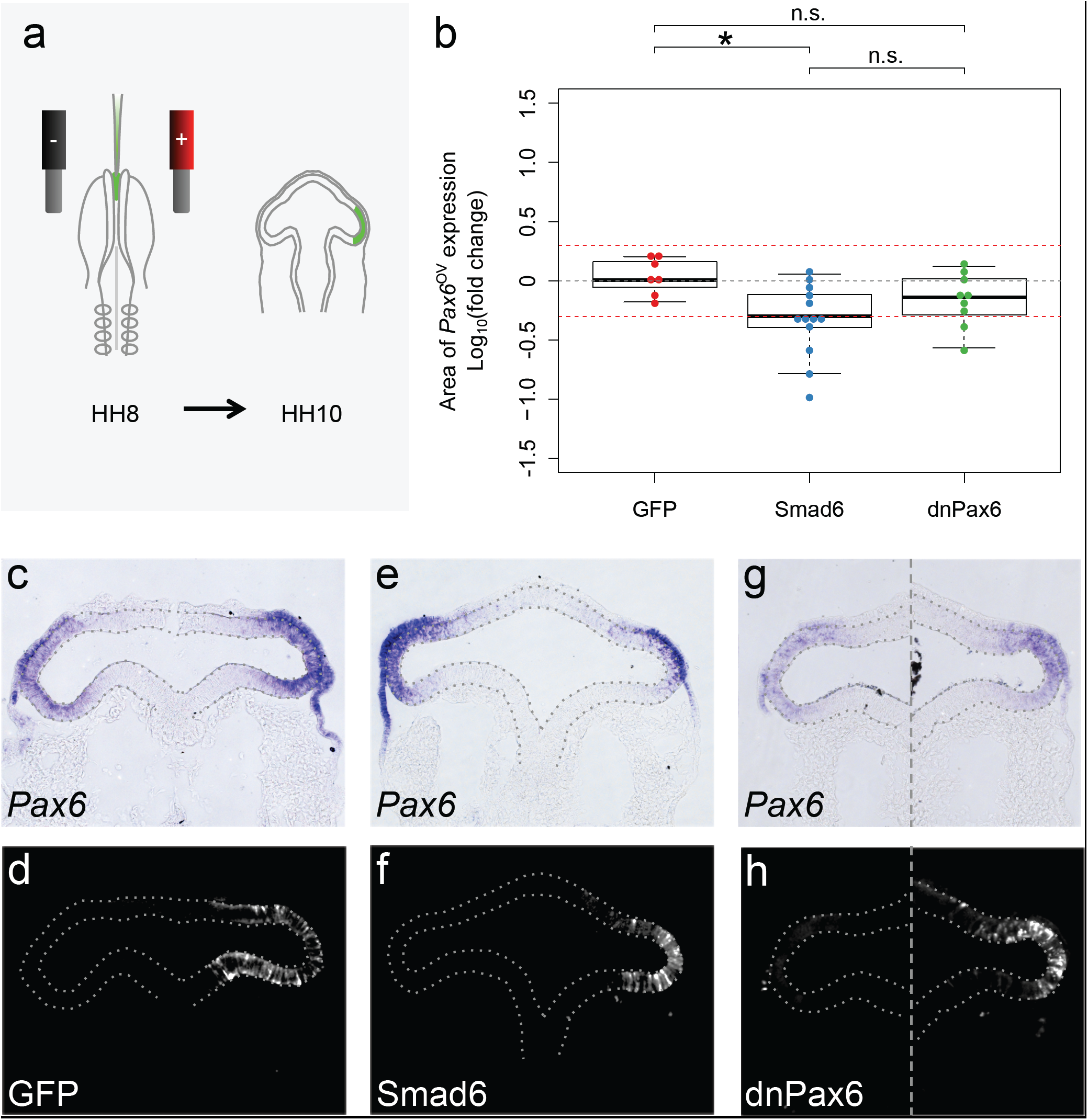
Bmp signalling is required for *Pax6* gene expression in the distal optic vesicle. **a)** DNA expression constructs were injected into the lumen of the anterior neural tube of stage HH8 chick embryos and electroporated to transfect a single prospective optic vesicle, the other serving as an un-transfected internal control. Embryos were cultured for 10-12 hours overnight until stage HH10 when they were analyzed. **b)** The sectional area of *Pax6* gene expression was measured and compared between electroporated an non-electroporated optic vesicles for each embryo. Log10(fold change) was plotted for embryos electroporated with GFP control construct, Smad6 + GFP construct or dnPax6 + GFP construct. Red guide lines indicate the level of +/-2-fold change in sectional expression area. * P < 0.05; n.s. indicates P >= 0.05. **c)** *Pax6* gene expression following transfection with GFP control. **d)** anti-GFP immunofluorescence showing location of GFP transfected cells. **e)** *Pax6* expression following transfection with Smad6 + GFP. **f)** anti-GFP immuno showing location of Smad6 + GFP transfected cells. **g)** *Pax6* expression following transfection with dnPax6 + GFP. **h)** anti-GFP immuno showing location of dnPax6 + GFP transfected cells. Optic vesicles are indicated by broken outlines.

Auto-regulation of *Pax6* has been reported in a number of tissues including the lens^17^. To test for *Pax6* auto-regulation in the optic vesicle, a C-terminally truncated dominant negative *Pax6* gene (*dnPax6*)^18^ was mis-expressed unilaterally, while a C-terminal riboprobe was used to selectively detect endogenous *Pax6* expression. *dnPax6* did not disrupt endogenous *Pax6* expression (0.75 ± 0.36 fold; *P* > 0.05; *n* = 9; Fig. 2b, g, h) compared with the GFP control, yet nor could we distinguish a difference between *dnPax6* and *Smad6* mis-expression (Fig. 2b; *P*> 0.05). Thus, while distal *Pax6* expression in the optic vesicle requires Bmp signalling *in vivo*, we cannot exclude the possibility that upstream Bmp action may mask subsequent *Pax6* auto-regulation.

Migratory neural crest cells reach the optic vesicle at stage HH10 and are thought to induce proximal and suppress distal gene expression via Tgfb subfamily signalling^19, 20^. Exogenously supplied Tgfb subfamily ligand (Activin A) was reported to induce proximal (*Wnt2b, Mitf*) and inhibit distal (*Pax6, Vsx2*) gene expression in explant cultures^19^. In contrast to this tissue-extrinsic induction mechanism, stem cell-derived retinal organoids are reported to polarise tissue-autonomously, exemplified by the spontaneous acquisition of proximal Wnt activity^21^. This raises the possibility of a redundant tissue-intrinsic polarising activity. Given that distal *Tgfb2* expression correlates with *Pax6* (Fig. 1a & e) we asked whether *Pax6* might induce *Tgfb2* to activate proximal target genes tissue-autonomously. In comparison with GFP controls (1.06 ± 0.17 fold; *n* = 8; Fig. 3a, b, c), mis-expression of *dnPax6* in single optic vesicles diminished *Tgfb2* expression relative to contralateral control vesicles (0.79 ± 0.54 fold; *P* < 0.05; *n* = 15; Fig. 3a, d, e). Thus, the *Pax6* master controller is required for *Tgfb2* expression in the distal vesicle, consistent with a report of Pax6 binding sites located within the *Tgfb2* promoter^22^.

**Figure 3:**
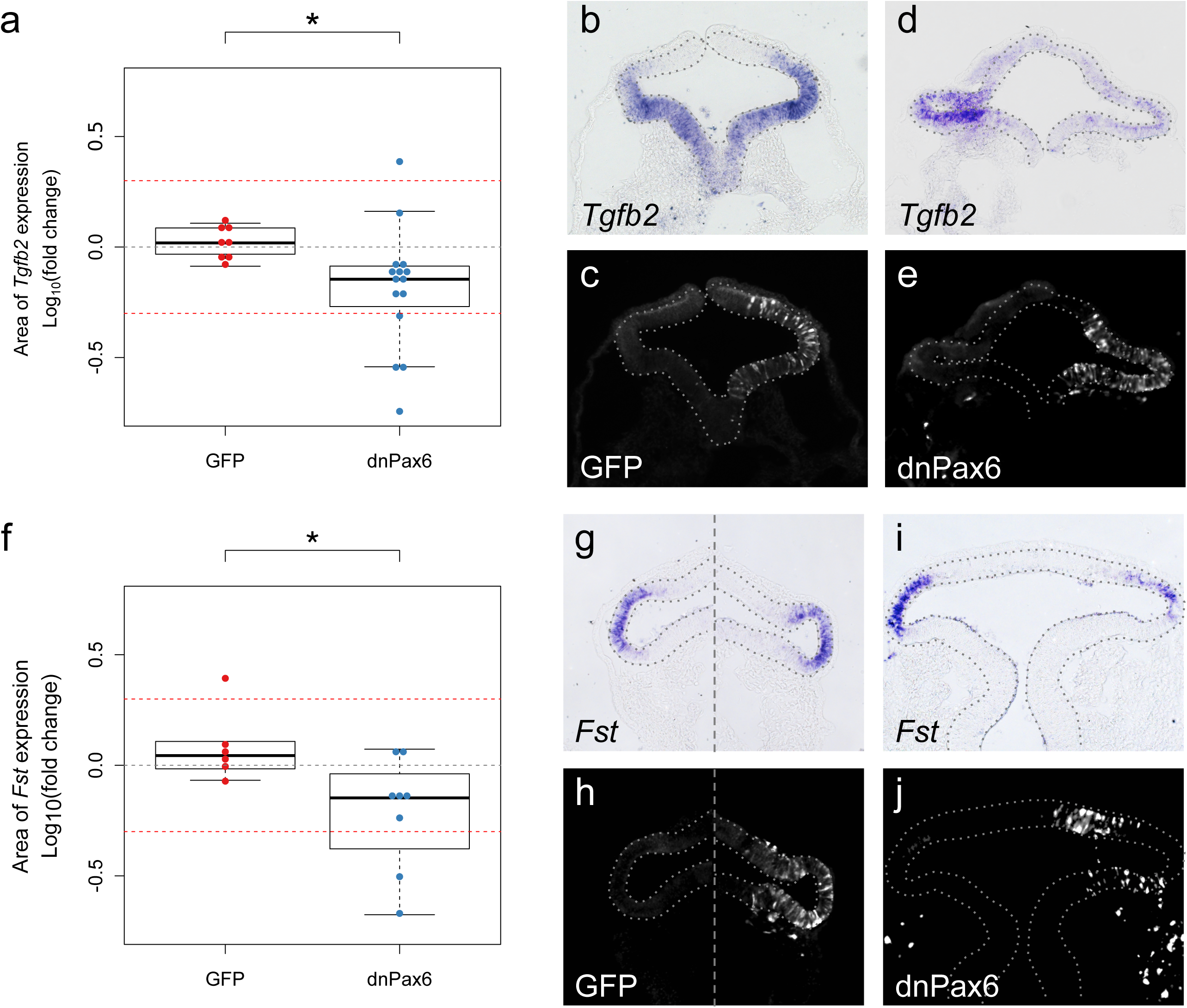
Pax6 function is required for expression of *Tgfb2* and *Fst*. **a-e)** *Tgfb2* gene expression was assessed 12 hours after electroporation of GFP or dnPax6 + GFP into a single optic vesicle. **a)** Sectional area of *Tgfb2* gene expression was measured and compared between electroporated an non-electroporated optic vesicles for each embryo. Log10(fold change) was plotted for each embryo. Red guide lines indicate the level of +/-2-fold change in sectional expression area. **b)** *Tgfb2* gene expression following electroporation with GFP control. **c)** anti-GFP immunofluorescence showing location of GFP transfected cells. **d)** *Tgfb2* gene expression following electroporation with dnPax6 + GFP. **e)** anti-GFP immuno showing location of dnPax6 + GFP transfected cells. **f-j)** *Fst* expression was assessed 12 hours after electroporation with GFP or dnPax6 + GFP. **f)** Sectional area of *Fst* gene expression was measured and compared between electroporated an non-electroporated optic vesicles for each embryo. Log10(fold change) was plotted for each embryo. Red guide lines indicate the level of +/-2-fold change in sectional expression area. **g)** *Fst* gene expression following electroporation with GFP control. **h)** anti-GFP immuno showing location of GFP transfected cells. **i)** *Fst* expression following electroporation with dnPax6 + GFP. **j)** anti-GFP immuno showing location of dnPax6 + GFP transfected cells. Optic vesicles are indicated by broken outlines. * P < 0.05.

This presents a paradox however; *Tgfb2* expression (Fig. 1e) negatively correlates with its positive targets *Wnt2b* and *Mitf* (Fig. 1c, d), yet positively correlates with its negative targets *Pax6* and *Vsx2* (Fig. 1a, b)^19^. How might Tgfb pathway activation become inverted relative to *Tgfb2* gene expression? We considered whether *Pax6* might also activate *Fst* (Fig. 1f), a Tgfb antagonist, to grant distal immunity from Tgfb signalling. Compared with GFP controls (1.31 ± 0.63 fold; *n* = 6; Fig. 3f, g, h), mis-expression of *dnPax6* in a single optic vesicle significantly reduced *Fst* expression (0.69 ± 0.34 fold; *P* < 0.05; *n* = 8; Fig. 3f, i, j). Thus, *Pax6* function is additionally required for *Fst* expression in the distal vesicle.

The paradoxical out-of-phase expression of distal *Tgfb2* and its proximal (positive) targets might then be explained by differential diffusion of *Tgfb2* and *Fst* gene products resulting in: i) Tgfb2 being locally sequestered by slow-diffusing Fst within the distal vesicle, thereby preserving distal character; ii) fast-diffusing Tgfb2 dispersing proximally away from Fst, to induce proximal character within the neighboring proximal vesicle.

To test if this hypothesis is plausible, we examined a reaction-diffusion model of the interactions summarised in Fig. 4a (Model A; see Supplementary Information) and performed numerical simulations with a variety of diffusion ratios for Tgfb2 dimers and Fst monomers versus Fst:Tgfb2 complexes (e.g. Fig. 4b, c; Supplementary Movie 1). Simulations demonstrated that local inhibition and lateral-activation of Tgfb signalling may occur if the diffusion rate of Fst:Tgfb2 complexes exceed that of Fst monomers. Although initially counter-intuitive, there is precedent for ligand:antagonist complexes that disperse faster than their individual constituents^23^ and our subsequent simulations assume this condition is satisfied.

**Figure 4:**
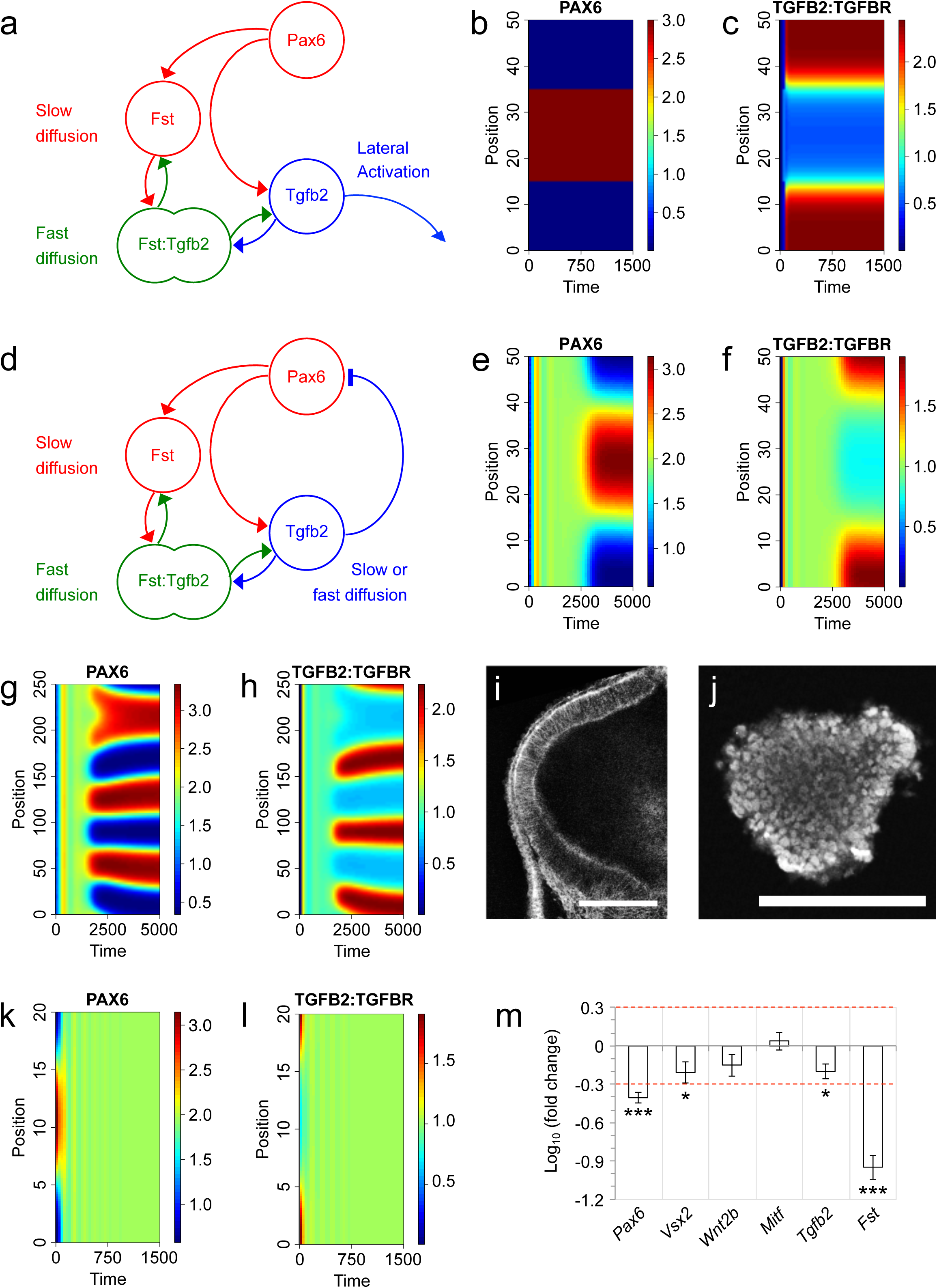
1-D Reaction-diffusion modelling of the *Pax6*/*Fst*/*Tgfb2* gene network. **a)** Summary of Model A in which Pax6 drives expression of both Fst and Tgfb2, whereas Fst inhibits Tgfb2 function via sequestration. Slow diffusion of Fst was postulated to result in local inhibition of Tgfb2 at the source of *Pax6*/*Tgfb2*/*Fst* expression. Conversely, fast diffusion of Tgfb2 was postulated to drive lateral activation of its downstream signalling pathway away from the *Pax6*/*Fst*/*Tgfb2*-expressing region. **b-c)** Numerical simulation of Model A in which Pax6 expression is regionally restricted throughout. For all simulations, units of space, time and molecular concentrations are arbitrary. The graphs depict the time-evolution (x-axis) for 1-D spatial distributions (y-axis) of: **b)** Pax6, and **c)** activated Tgfb2:Tgfb-receptor signalling complex. **d)** Summary of Model B in which Fst:Tgfb2 complex quickly diffuses and dissociates while Tgfb2 additionally inhibits Pax6 transcriptional activator function. **e-f)** Numerical simulation of Model B in which Pax6 expression is initially homogenous but noisy. The graphs depict spontaneous generation of **e)** a Pax6+ ‘distal pole’ flanked by **f)** Tgfb2:Tgfbr+ ‘proximal poles’. **g-h)** The Model B simulation of e-f was repeated with a larger tissue size yielding multiple Pax6+ ‘distal poles’ interspersed with Tgfb2:Tgfbr+ ‘proximal poles’. **i)** Confocal section of an HH10 tg(membrane-GFP) embryo showing optic vesicle size prior to explant culture. **j)** Confocal section of a fixed optic vesicle explant showing the collapsed tissue following 16 hours culture. Cell nuclei are stained with Propidium Iodide. **k-l)** Continuation of the Model B simulation shown in e-f, in which the previously patterned tissue was reduced in size by 0.4-fold. The simulation predicts a rapid de-polarisation of both **k)** Pax6 and **l)** active Tgfb2:Tgfbr signalling complex. **m)** RT-QPCR analysis of proximal and distal marker gene expression in response to optic vesicle collapse during 16-hour explant culture. Values plotted are Log10(mean fold change) +/-SEM. Red guide lines indicate the levels of +/-2-fold change in gene expression. * P < 0.05; ** P < 0.01; *** P < 0.001.

Given that Tgfb signalling is known to disrupt Pax6 protein function^18^, such local inhibition and lateral-activation of Tgfb signalling equates to local positive feedback and lateral-inhibition of the *Pax6* master control gene, respectively (Fig. 4d). This is functionally equivalent to a simple Activator-Inhibitor^24^ type Turing network^7^, which can serve as a spontaneous pattern generator; *Pax6* and *Fst* comprising a short-range auto-regulating Activator, and *Tgfb2* as the long-range Inhibitor. To explore whether the network of Fig. 4d possesses spontaneous polarising activity, we simply extended Model A to include inhibition of Pax6 function by Tgfb signalling (Model B; see Supplementary Information). Simulations showed that an initially homogenous but noisy *Pax6* distribution is readily converted into a polarised pattern, wherein *Pax6* expression becomes regionally restricted (Fig. 4e) and out-of-phase with Tgfb receptor activation (Fig. 4f; Supplementary Movie 2). Additionally, simulating larger tissue sizes results not in a larger *Pax6*-expressing distal pole, but in a greater number of *Pax6*-expressing distal poles of approximately equal size (Fig. 4g, h). This hallmark feature of Turing networks is remarkably consistent with observations of retinal organoid cultures in which stem cell aggregates yielded between one and four retinas each^8^.

It follows that reducing tissue size should limit the number of *Pax6*-expressing distal poles until polarisation is no longer possible. When cultured as isolated explants in the absence of serum, polarised HH10 optic vesicles (e.g. Fig. 4i) collapse into compact spheroids (Fig. 4j) reducing this tissue’s longest dimension to ≤0.5 fold. Continuing the Model B simulation of Fig. 4e & f after reducing the length of this polarised tissue predicts that proximal-distal polarity should be lost in this case (Fig. 4k, l). To test this, we compared proximal and distal gene expression before and after vesicle collapse under neutral culture conditions. In agreement with the simulation, the four distal markers *Pax6* (0.40 ± 0.08 fold; *P* < 0.001; *n* = 4), *Fst* (0.12 ± 0.04 fold; *P* < 0.001; *n* = 4), *Tgfb2* (0.65 ± 0.17 fold; *P* < 0.05; *n* = 4) and *Vsx2* (0.66 ± 0.25 fold; *P* < 0.05; *n* = 4) are all reduced following vesicle collapse (Fig. 4m).

Model B (Fig. 4d) further predicts that interference with *Fst* gene expression should derepress Tgfb signalling and inhibit Pax6 protein function in the distal vesicle, via the direct Tgfb-dependent interaction of Smad3 with Pax6^18^. Moreover, if Pax6 auto-regulates in the distal vesicle, this should manifest as a Tgfb-mediated reduction in *Pax6* gene expression. To test this prediction, we employed morpholino oligonucleotides to suppress Fst translation within single optic vesicles and compared *Pax6* expression between these and unperturbed contralateral vesicles. Fst morpholino (FstMO) was first shown by Western Blotting to suppress endogenous Fst protein expression in cultured chick embryonic cells, as compared to a standard control morpholino (StdMO) that does not target *Fst* (Fig. 5a). *In vivo*, StdMO controls had no impact on *Pax6* expression in transfected optic vesicles (1.05 ± 0.31 fold; *n* = 20; Fig. 6b, c, d). In comparison, FstMO reduced *Pax6* expression in transfected vesicles (0.76 ± 0.50 fold; *P* < 0.01; *n* = 18; Fig. 5b, e, f), as predicted by Model B. We were able to rescue this loss of *Pax6* expression by co-transfecting FstMO together with an exogenous *Fst* transgene that evades FstMO (0.98 ± 0.35 fold; *P* > 0.05; *n* = 25; Fig. 5b, g, h), confirming that loss of *Pax6* was not due to a morpholino off-target effect. Thus, *Fst* is required for distal *Pax6* expression in the optic vesicle. This is consistent with earlier reports that neural induction by way of *Fst* overexpression induces *Pax6* in *Xenopus* animal cap explants^25^.

**Figure 5:**
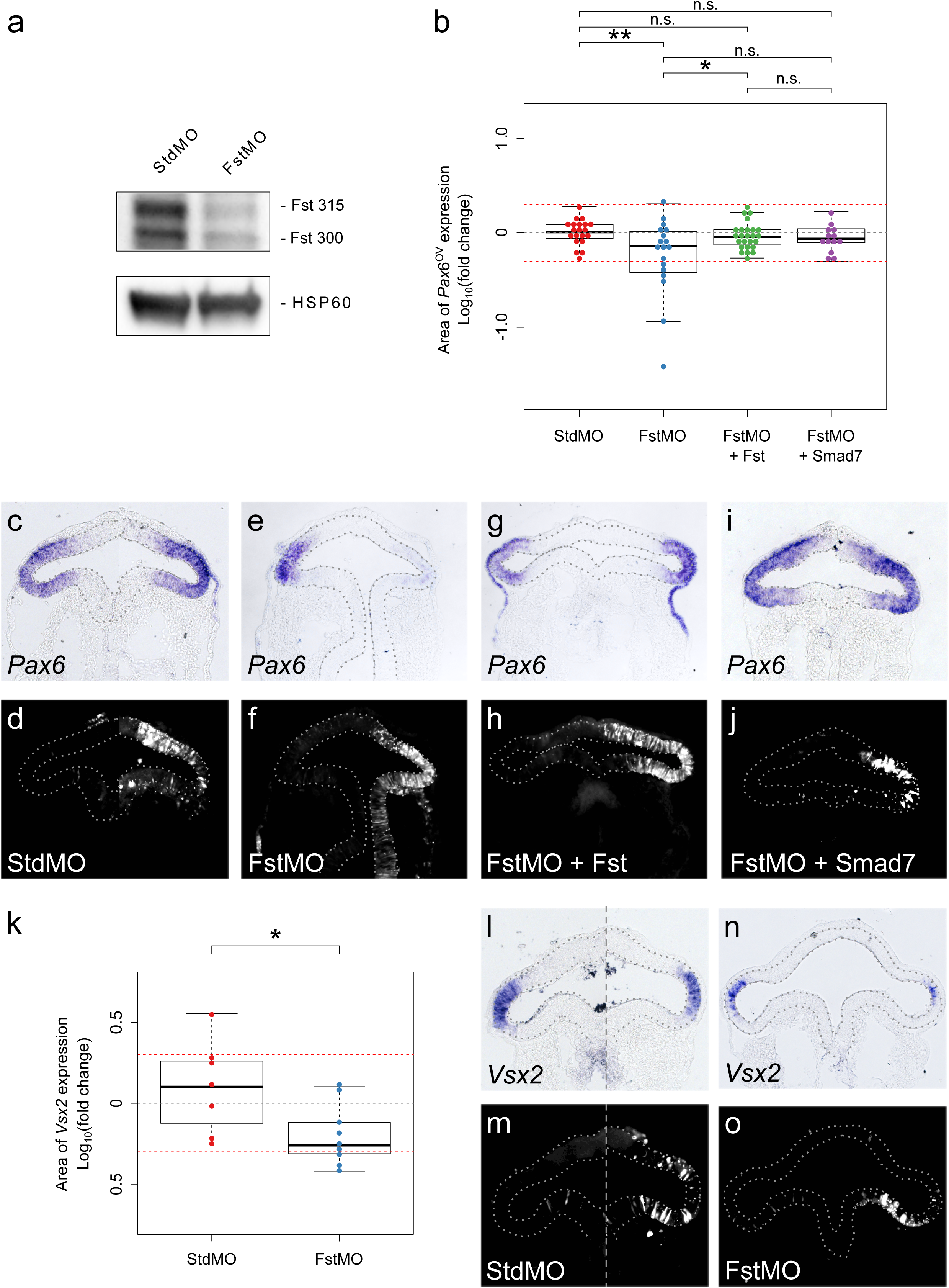
*Fst* gene function is required for correct optic vesicle polarisation via distal inhibition of Tgfb signalling. **a)** Western blot validation of Fst protein knockdown by FstMO but not StdMO in cultured chick embryo cells. **b-j)** Sectional area of *Pax6* gene expression was assessed 12 hours after co-electroporation of single optic vesicles with control/experimental morpholinos plus various gene expression constructs. **b)** Sectional area of *Pax6* gene expression was measured and compared between electroporated an non-electroporated optic vesicles for each embryo. Log10(fold change) was plotted for each embryo. Red guide lines indicate the level of +/-2-fold change in sectional expression area. **c)** *Pax6* gene expression following co-electroporation of standard control morpholino (StdMO) + GFP. **d)** FITC-labelled StdMO fluorescence showing location of transfected cells in panel c. **e)** *Pax6* gene expression following co-electroporation of *Fst* morpholino (FstMO) + GFP. **f)** FITC-labelled FstMO fluorescence showing location of transfected cells in panel e. **g)** *Pax6* gene expression following co-electroporation of FstMO + Fst gene expression construct. **h)** FITC-labelled FstMO fluorescence showing location of transfected cells in panel g. **i)** *Pax6* gene expression following co-electroporation of FstMO + Smad7 gene expression construct. **j)** FITC-labelled FstMO fluorescence showing location of transfected cells in panel i. **k-o)** Sectional area of *Vsx2* gene expression was assessed 12 hours after co-electroporation of single optic vesicles with control/experimental morpholino. **k)** Sectional area of *Vsx2* gene expression was measured and compared between electroporated an non-electroporated optic vesicles for each embryo. Log10(fold change) was plotted for each embryo. Red guide lines indicate the level of +/-2-fold change in sectional expression area. **l)** *Vsx2* gene expression following co-electroporation of StdMO + GFP. **m)** FITC-labelled StdMO fluorescence showing location of transfected cells in panel l. **n)** *Vsx2* gene expression following co-electroporation of FstMO + GFP. **o)** FITC-labelled FstMO fluorescence showing location of transfected cells in panel l. Optic vesicles are indicated by broken outlines. * P < 0.05; ** P < 0.01.

**Figure 6:**
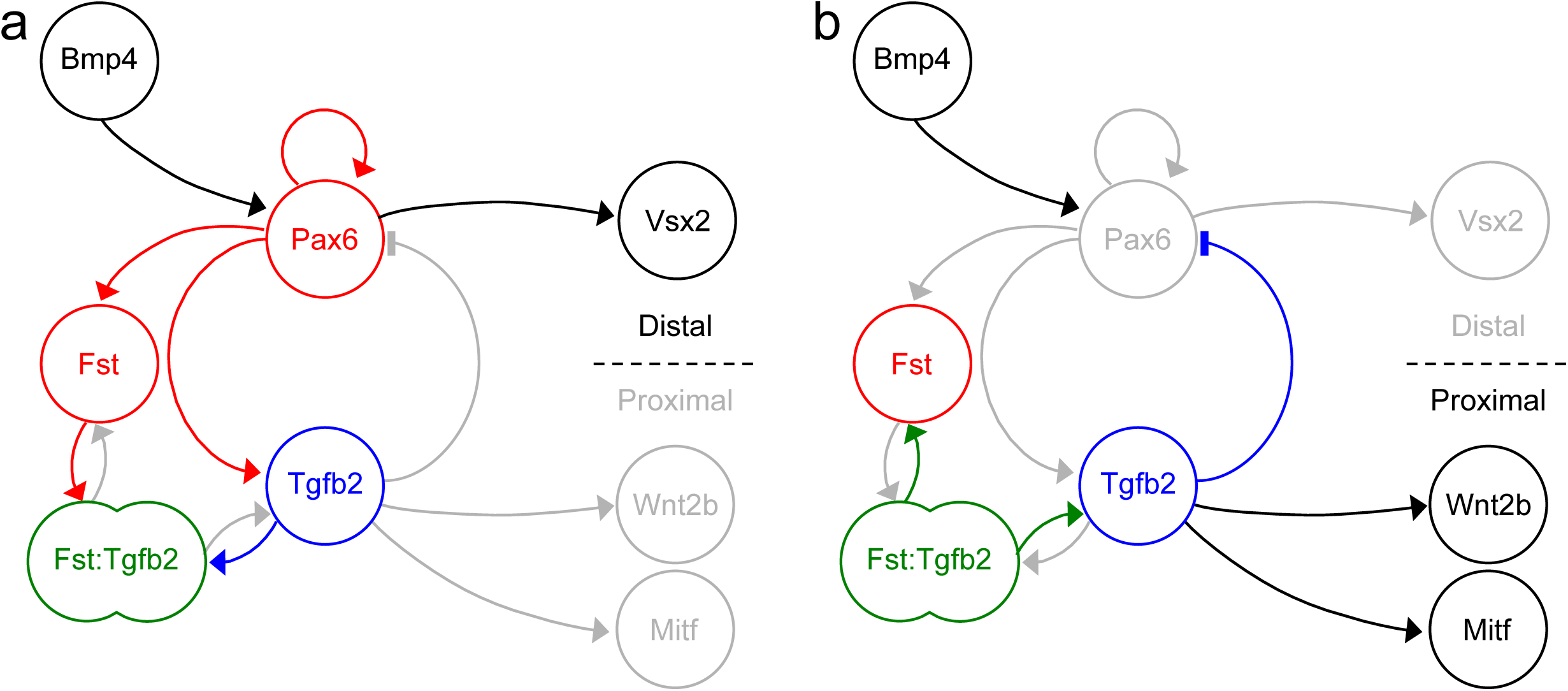
Proposed *Pax6*/*Fst*/*Tgfb2* network function during optic vesicle polarisation *in vivo*. **a)** At the prospective distal pole *Pax6* expression is promoted by upstream Bmp and reinforced via autoregulation. Pax6 drives distal expression of *Fst, Tgfb2* and downstream *Vsx2*. A molar excess of slow-diffusing Fst over Tgfb receptors is postulated to reversibly sequester Tgfb2 into fast-diffusing Fst:Tgfb2 complexes. **b)** At the prospective proximal vesicle, dissociation of fast-diffusing Fst:Tgfb2 complexes is postulated to release Tgfb2. A molar excess of Tgfb receptors over slow-diffusing Fst then permits receptor activation by Tgfb2, causing functional inhibition of Pax6 and induction of proximal markers *Wnt2b* and *Mitf*.

To verify that loss of *Pax6* expression is indeed due to the predicted de-repression of Tgfb signalling, we attempted an alternate rescue by co-transfecting FstMO together with a cell-autonomous Tgfb pathway inhibitor, *Smad7*. As can be seen (Fig. 5b, i, j), no significant loss of *Pax6* expression was observed (0.91 ± 0.31 fold; *P* > 0.05; *n* = 13) when Fst translation and Tgfb signalling were simultaneously suppressed.

In addition to inducing *Pax6*^25^, overexpression of *Fst* in *Xenopus* animal cap explants was reported to induce expression of the retinal photoreceptor marker *Opsin*^26^. We therefore investigated whether *Vsx2*, a distally expressed neural retinal marker^27^ (Fig. 1b), is similarly affected upon disruption of the *Pax6*/*Fst*/*Tgfb2* gene network. In comparison to StdMO controls (1.51 ± 1.05 fold; *n* = 7; Fig. 5k, l, m), FstMO significantly reduced distal *Vsx2* expression in transfected optic vesicles (0.69 ± 0.33 fold; *P* < 0.05; *n* = 9; Fig. 5k, n, o). Thus, de-repression of endogenous Tgfb signalling in the distal vesicle is detrimental for correct proximal-distal patterning, including specification of the neural retina. These results are consistent with Model B (Fig. 4d) and support the idea that Fst and Tgfb2 morphogens positively and negatively regulate Pax6 function, respectively, in order to polarise the optic vesicle.

## Discussion

The question of *Pax6*’s master control mechanism has been unresolved for a quarter of a century^28^. Here we have shown that the vertebrate *Pax6* directs expression of a pair of morphogen coding genes, *Fst* and *Tgfb2*, which modulate Pax6 function via positive and negative feedbacks. Our reaction-diffusion simulations showed that the *Pax6*/*Fst*/*Tgfb2* gene network may act as a self-organising Turing network, providing certain assumptions are satisfied. For instance, we have assumed that larger Fst:Tgfb2 complexes diffuse more quickly than smaller Fst monomers. This is counter-intuitive since pure diffusion rate is a function of molecular mass, yet there is precedent for this phenomenon; e.g. Sfrp:Wnt complexes have been observed to diffuse further than Wnt alone^23^. We postulate that Fst monomers disperse sub-diffusively due to binding interactions with extra-cellular matrix components and/or cell surface factors, e.g. heparin sulfate proteoglycans^29^. In the context of Fst:Tgfb2 complexes, interaction surfaces may be shielded enabling the larger complex to disperse further and faster than its constituents.

The assumed rapid dispersal of Fst:Tgfb2 complexes is only required if Tgfb2 sequestration by Fst is reversible, which is currently unknown. Low affinity Fst:Bmp interactions are known to be reversible whereas high affinity Fst:Activin interactions are effectively irreversible^30^. If Fst:Tgfb2 associate irreversibly then spontaneous pattern formation is still possible, but it changes assumptions regarding effective diffusion rates: Fst:Tgfb2 diffusion would then become irrelevant and instead, Tgfb2 dimers must diffuse faster than Fst monomers^31^.

By demonstrating how *Pax6* may drive self-organisation of the primary retinal axis, our findings offer the first mechanistic explanation of *Pax6*‘s long-known but poorly understood master control function. In the embryo, we propose that this putative Turing network acts to self-organise the optic vesicle’s proximal-distal axis (as summarised in Fig. 6a & b) and that previously identified inductive interactions serve to trigger and/or synchronise this tissue-autonomous activity with neighboring tissues. For instance, Bmp signals from the overlying head ectoderm may bias proximal-distal polarity to align the distal pole with the prospective lens. BMP inhibition^14^ (or an absence of lens ectoderm) may then prevent the reported *Pax6*/*Fst*/*Tgfb2* network from being activated, or else permit supplementary Tgfbs from the surrounding neural crest mesenchyme^19, 20^ to overwhelm this tissue-intrinsic polarising activity. In turn, it has not escaped our attention that distal *Fst* may mediate classical lens induction^9^ by opposing these same lens-inhibitory Tgfb signals^20^; indeed, *Fst* overexpression induces lens crystallin expression in *Xenopus* animal cap explants^25^.

We did not investigate the role of Wnt in establishing proximal identity within the optic vesicle, except to test for direct synergism between Wnt and Bmp as previously proposed^15^. In the absence of such synergism, we suggest that Wnt acts downstream of the *Pax6*/*Fst*/*Tgfb2* gene network, since i) *Wnt2b* is a Tgfb target gene^19^ restricted to the proximal optic vesicle (Fig. 1c), and ii) expression of *Wnt2b* is absent from the peri-ocular surface ectoderm until HH11^20^ prior to which, polarised *Wnt2b* expression is already established within the optic vesicle itself (Fig. 1c).

During retinal organoid development *in vitro*^8^, we propose that the *Pax6*/*Fst*/*Tgfb2* network may suffice to self-organise the retina’s primary axis in the absence of well-ordered tissue interactions characteristic of embryonic eye development. The comparatively chaotic nature of organoids makes them an ideal counterpart to embryonic models of development as they can unmask cryptic self-organising mechanisms and test them to breaking point; contrast the straightforward elaboration of an existing pre-pattern (Fig. 4b, c; analogous to localised *Pax6* induction by neighboring Bmps *in vivo*) with the more turbulent emergence of order from disorder (Fig. 4e, f; analogous to spontaneous *Pax6* activation in retinal organoids). Further exploration of the *Pax6*/*Fst*/*Tgfb2* network may drive future developments in retinal organoid technology and help underpin applications in disease modelling, drug discovery and regenerative therapies. Given the deeply homologous nature of *Pax6*’s master control function, we would predict that *Pax6* orthologues participate in functionally homologous Turing networks in non-vertebrates, which may comprise the same or different morphogens.

## Materials & Methods

### Chick embryos

Fertile brown hen’s eggs (Henry Stewart) were incubated at 38 °C in a humidified incubator until the required stage of development: HH8 for *in ovo* electroporation experiments; HH10 for *in vitro* explant experiments.

### Explant Assays

HH10 embryos were incubated with 0.25 % Trypsin-EDTA at 38 °C for 7 minutes. Trypsin was then de-activated by transferring into 20 % chick serum on ice for 5 minutes. Embryos were then washed with Tyrodes solution and pinned onto Sylgard-coated dissection dishes. Head surface ectoderm and peri-ocular mesenchyme were carefully removed using 30 gauge syringe needles from both dorsal and ventral sides. Once cleaned, both optic vesicles were removed and held in Tyrodes solution on ice. Left and right optic vesicles were separately pooled from at least five embryos, yielding two match-paired pools for use as treated and control samples. Pooled vesicles were cultured in polyHEMA (Sigma) coated culture wells to prevent adhesion, with DMEM-F12 media (Invitrogen) supplemented with 1X N2 (Invitrogen), 1X L-Glutamate and 1X Penicillin/Streptomycin at 37 °C and 5% CO_2_ for 16 hrs. Culture media for treated samples was supplemented with the following factors as required: 35 ng/ml Bmp4 (R&D Systems), 0.5 μM BIO (Sigma) with 0.1 % DMSO (Sigma), or 0.1 % DMSO only.

### Quantitative RT-PCR

Explant samples were lysed in 1 ml Trizol (Ambion) and processed for total RNA extraction. RNA samples were digested with DNase I (Ambion) and re-extracted by acidic Phenol/Chloroform. RNA concentrations were determined by NanoDrop ND-1000 Spectrophotometer. For each experiment, equal quantities of treated and control sample RNA (typically between 0.1 – 0.6 μg) were used as template for first strand cDNA synthesis using Superscript II reverse transcriptase (Invitrogen) and random hexamers. cDNAs were diluted 1:20 before relative quantitation of transcript levels by real-time PCR using SYBR Green master mix (Applied Biosystems) and target-specific primers (Supplementary Table 1). Relative transcript quantification was via the standard curve method, and target gene expression was normalised to the reference gene β-Actin. Fold changes were calculated for each matched-pair (treated/control) then log-transformed to ensure normal distribution. These were plotted as mean +/-standard deviation. Student’s paired t-test was used to calculate the probability of the observed (or more extreme) differences between match-paired (treated and control) sample means assuming that the null hypothesis is true.

### Morpholino Knockdown Validation

*Fst*-expressing somite tissue from wild-type chick embryos were dissected and cultured in Dulbecco’s Modified Eagle Medium, 10% foetal bovine serum, 1% penicillin/streptomycin for 4 h before transfecting with 1 mM translation-blocking FstMO (Gene Tools; sequence 5’-GATCCTCTGATTTAACATCCTCAGC-3’) or 1mM StdMO negative control (Gene Tools; sequence 5’-CCTCTTACCTCAGTTACAATTTATA-3’) using Endoporter PEG (Gene Tools). Protein was extracted after 48 h. Protein lysate (30 μg) was run on pre-cast 4-15% polyacrylamide gels (Bio-Rad) and blotted onto polyvinylidene fluoride membrane (Bio-Rad). Primary antibody against Fst (Abcam ab47941; 1:2,000) was applied at 4°C overnight and secondary polyclonal goat anti-rabbit-HRP (Cell Signaling Technology #7074; 1:2,000) was applied for 1 h at room temperature. Primary antibody against HSC70 (Santa Cruz sc-7298; 1:2,500) was applied at 4°C overnight and secondary polyclonal goat anti-mouse-HRP (Agilent P0447; 1:1,000) was applied for 1 h at room temperature. The blots were treated with an ECL substrate kit and imaged.

### *In Ovo* Embryo Electroporation

Plasmid DNA (2 – 5 μg/ul) or plasmid DNA and FITC-labelled Morpholino oligonucleotides (2 μg/ul and 0.5 mM, respectively), were injected into the open neural tube of stage HH8 chick embryos *in ovo* (Fig. 2a). A pair of platinum electrodes connected to an Ovodyne electroporator and current amplifier (Intracel) were then used to electroporate the DNA or DNA + Morpholino into either left or right side of the anterior neural tube via 4 pulses of 22 volts with 50 ms duration and at 1 second intervals. Once electroporated, embryos were sealed with adhesive tape and incubated for 10 – 12 hours at 38 °C until embryos had reached stage HH10.

### Wholemount In Situ Hybridization and Immunofluorescence

Embryos were fixed in 4% PFA overnight at 4 °C, then dehydrated by methanol series and stored at −20 °C. Following rehydration, embryos were processed for wholemount in situ hybridization using 1 μg/ml DIG-labelled antisense probes for *Pax6, Vsx2, Mitf, Fst* (see Supplementary Table 2 for PCR primers), *Tgfb2* (EST clone ChEST262a17)^32^, *Wnt2b* (a gift from Susan Chapman) and *Bmp4* (a gift from Elisa Martí). Probes were hybridized at 65 °C for up to 72 hrs. After incubation with 1:5,000 anti-DIG antibody (Roche) and washing, 4.5 μl nitroblue tetrazolium (50 mg/ml) and 3.5 μl 5-bromo-4-chloro-3-indolyl phosphate (50 mg/ml) per 1.5 ml developing solution were used for color development. Embryos were embedded in 7.5 % gelatin, 15 % sucrose and cryo-sectioned at 15 μm thickness. Following de-gelatinisation, sections were blocked in PBTS buffer (PBS with 2 % BSA, 0.1 % Triton X-100 and 10 % goat serum) for 1 hr at room temperature. EGFP transgene expression was then detected using rabbit anti-GFP primary antibody (Abcam; 1:500 dilution) and Alexa568 goat anti-rabbit secondary antibody (Invitrogen; 1:1000 dilution). Morpholino FITC fluorescence was observed directly. Labelled sections were imaged using a 20X objective on an Axioplan widefield fluorescence microscope with Axiocam HRc camera and Axiovision software (Carl Zeiss).

### Relative Quantification of In Situ Hybridization Staining

Assuming that average cell size is invariant between left and right optic vesicles of the same embryo, then the relative area of staining is proportional to the relative number of cells exceeding a common detection threshold. To quantify this, brightfield micrographs were converted to greyscale, inverted then thresholded and the area of optic vesicle staining measured in FIJI^33^. Transfected and contralateral controls from the same embryo were processed simultaneously to ensure identical treatment. Staining area in transfected vesicles was then normalised to internal contralateral controls, yielding fold change in gene expression area. Fold changes were log-transformed to ensure normal distribution prior to plotting and null hypothesis significance testing. Box plots showing mean Log10(fold change) +/-standard deviation were generated in R with the package ‘Beeswarm’. Welch’s two-sample t-test (for pairwise comparisons) or one-way ANOVA with Tukey’s post hoc test (for groupwise comparisons) were used to calculate the probability of the observed (or more extreme) differences between sample means assuming that the null hypothesis is true.

### Reaction-Diffusion Simulations

Partial differential equations were coded in R using the function tran.1d() from package ‘ReacTran’ to handle diffusion terms. 1-D numerical simulations used the function ode.1d() from package ‘deSolve’ and the default integrator. Parameter sweeps were performed to identify suitable diffusion rates (see Supplementary Movies 1 & 2). Simulations were run with both periodic and zero-flux boundary conditions, with comparable results. See Supplementary Information for model code and narrative text. The model code is explained in Supplementary Information, is available via our GitHub repository (https://github.com/GrocottLab/) and is accessible as an interactive Jupyter Notebook (https://mybinder.org/v2/gh/GrocottLab/Pax6-Fst-Tgfb2_Reaction_Diffusion_Models/master).

## Acknowledgements

This work was supported by a Fight for Sight UK Early Career Investigator Award to T.G. (1365/66), a BBSRC Project Grant (BB/N007034/1) to A.E.M. and a H2020 Marie Sklodowska-Curie Actions Individual Fellowship (705089) to E.L.-V. We thank Paul Thomas of the Henry Wellcome Laboratory for Cell Imaging for assistance with microscopy and colleagues in the laboratories of Grant Wheeler and Andrea Münsterberg for valuable discussions. We thank Elisa Martí and Susan Chapman for sharing plasmids.

## Author Contributions

T.G. conceived the project, designed/performed the experiments and computational modelling, analysed the data and prepared the figures. T.G. and A.E.M. interpreted the data and wrote the manuscript. G.F.M. and E.L.-V. performed morpholino knockdown validation.

---

**Supplementary Table 1.**
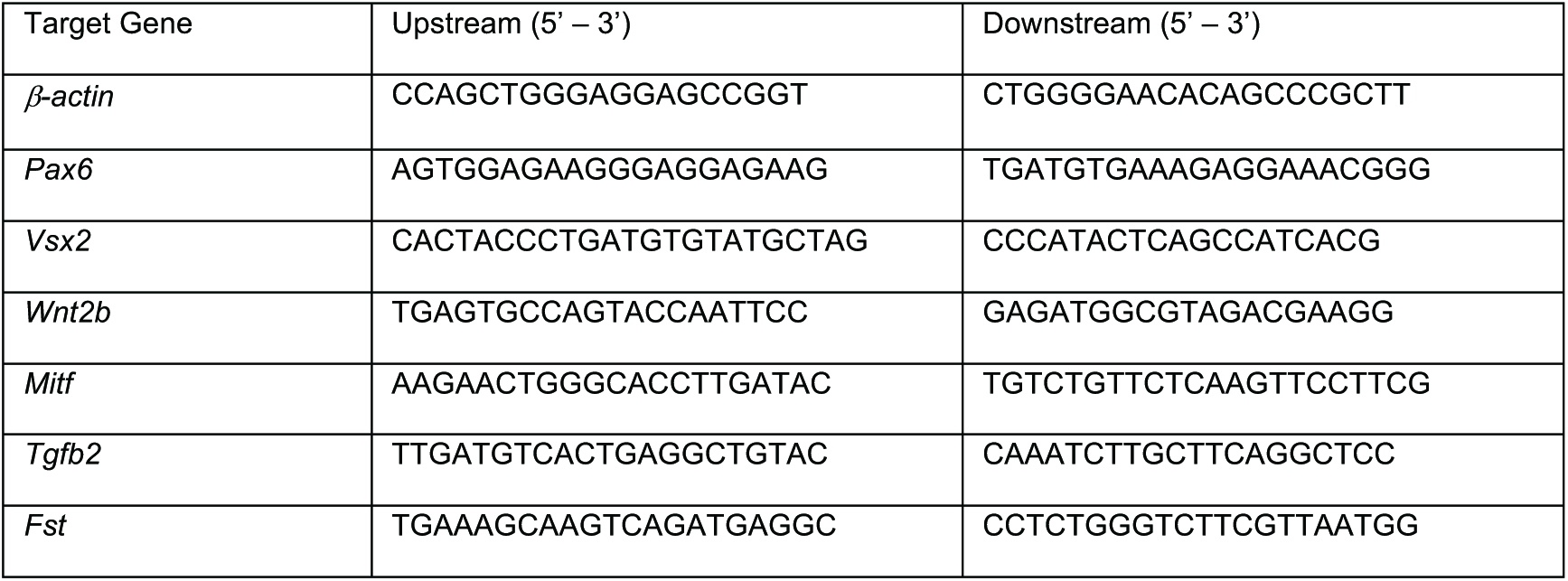
Primer sequences for qRT-PCR.

**Supplementary Table 2.**
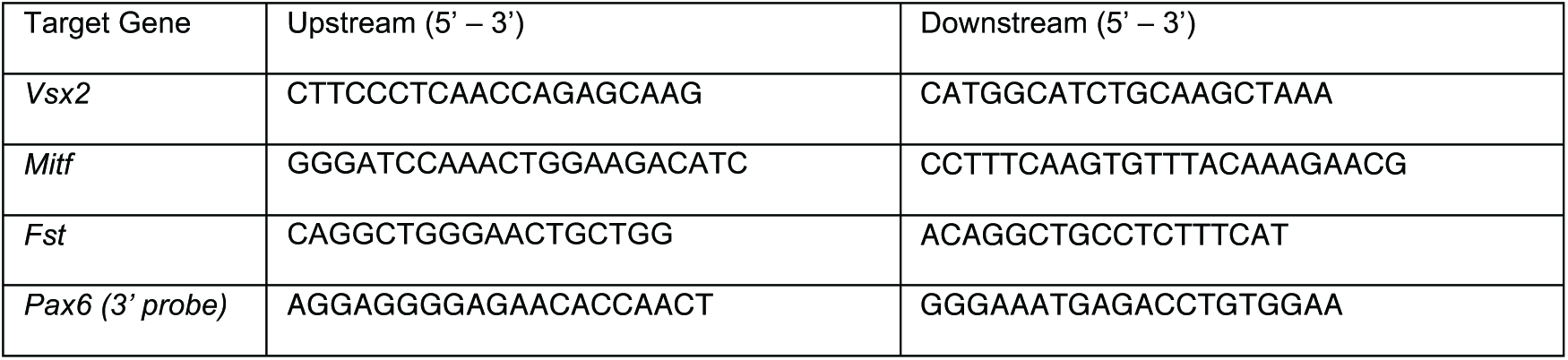
Primer sequences for PCR cloning of in situ hybridization probes.

