## Supplementary Information for "The *Pax6* master control gene initiates spontaneous retinal development via a self-organising Turing network"

#### Reaction-Diffusion Model A:

##### Introduction:

In this model, Pax6 drives expression of both Fst and Tgfb2. Fst monomers and Tgfb2 dimers may associate to form a hetero-tetrameric Fst:Tgfb2 complex. The Fst:Tgfb2 complex may dissociate in turn, thereby releasing Fst monomers and Tgfb2 dimers. Tgfb2 signals via association with Type I and Type II receptors, yielding an activated hetero-hexameric Tgfb2:Tgfbr signalling complex (comrpising one Tgfb2 ligand dimer and four Tgfb receptors - two Type I and two Type II; not shown here).

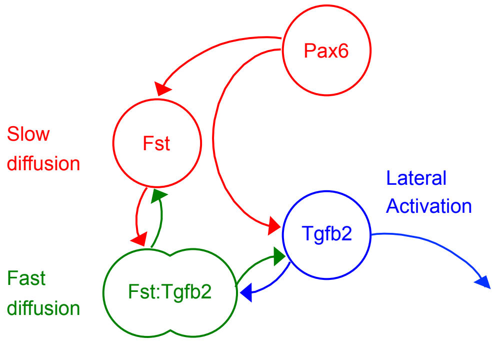


As dicussed in the main article, Fst monomers are assumed to exhibit slower effective diffusion than Fst:Tgfb2 complexes, whereas effective diffusion of Tgfb2 dimers relative to Fst:Tgfb2 complexes is less critical. If this differential diffusion condition is met, Model A is able to elaborate an existing Pax6 pre-pattern (e.g. expression induced within the distal optic vesicle by upstream BMP signalling) via 'lateral activation' of Tgfb receptor complexes away from the source of Pax6 expression (e.g. within the adjacent proximal optic vesicle). In other words, the pattern of Tgfb pathway activation becomes the inverse of upstream Pax6 expression.

##### Equations:

The following system of equations formally represents the above interactions. Each equation describes the rate of change of concentration for the various molecular species downstream of Pax6. The initial concentrations of Pax6 ($P$) and Tgfb receptors ($R$) are held constant throughout the simulation, i.e. their rates of change, $\frac{\partial[P]}{\partial t}$ and $\frac{\partial[R]}{\partial t}$ are set to zero throughout.

###### Fst:

$$\frac{\partial[F]}{\partial t} = a_F . \frac{[P]}{([P]+1)} - 2.a_{FT} . [F]^2 . [T] + 2.b_{FT} . [FT] - b_F . [F] + D_F \frac{d^2[F]}{dx^2}$$

Where,

- Pax6 ($P$) positively regulates Fst expression via Michaelis-Menton kinetics $\frac{[P]}{[P]+1}$.

- Pairs of Fst monomers are consumed in the formation of Fst:Tgfb2 complexes ($FT$) via $-2.a_{FT} . [F]^2 . [T]$, and released again via $+2.b_{FT} . [FT]$. This implements the two-way reaction: $1\,\mathrm{T} + 2\,\mathrm{F} \underset{b_{FT}}{\overset{a_{FT}}{\rightleftharpoons}} 1\,\mathrm{FT}$, whereby one Tgfb2 dimer reversibly associates with two Fst monomers.

- Fst is turned over via $-b_F . [F]$.

- Fst diffuses via $D_F \frac{d^2[F]}{dx^2}$.

##### Tgfb2:

$$\frac{\partial[T]}{\partial t} = \left(a_T.\frac{[P]}{([P]+1)}\right)^2 - a_{FT}.[F]^2.[T] + b_{FT}.[FT] - a_{RT}.[R]^4.[T] - b_T.[T] + D_T\frac{d^2[T]}{dx^2}$$

Where,

- Pax6 ($P$) positively regulates Tgfb2 expression via Michaelis-Menton kinetics $\frac{[P]}{[P]+1}$.

- Tgfb2 is produced/secreted as a disulphide linked dimer, hence $\left(a_T.\frac{[P]}{([P]+1)}\right)^2$.

- Tgfb2 is sequestered in the formation of Fst:Tgfb2 complexes ($FT$) via $-a_{FT}.[F]^2.[T]$, and released again via $+b_{FT}.[FT]$. This implements the two-way reaction: $1\,\mathrm{T} + 2\,\mathrm{F} \underset{b_{FT}}{\overset{a_{FT}}{\rightleftharpoons}} 1\,\mathrm{FT}$, whereby one Tgfb2 dimer reversibly associates with two Fst monomers.

- Tgfb2 is irreversibly consumed in the formation of activated receptor complexes ($RT$) via $-a_{RT}.[R]^4.[T]$. This implements the one-way reaction: $1\,\mathrm{T} + 4\,\mathrm{R} \overset{a_{RT}}{\longrightarrow} 1\,\mathrm{RT}$, whereby one Tgfb2 dimer associates with four Tgfbr monomers (two type I receptors + two type II receptors). This reaction is irreverible since cells internalise and ultimately turn over activated receptor complexes.

- Tgfb2 is turned over via $-b_T.[T]$.

- Tgfb2 diffuses via $D_T\frac{d^2[T]}{dx^2}$.

##### Fst:Tgfb2 complex:

$$\frac{\partial[FT]}{\partial t} = a_{FT}.[F]^2.[T] - b_{FT}.[FT] + D_{FT}\frac{d^2[FT]}{dx^2}$$

Where,

- A pair of Fst monomers and a single Tgfb2 dimer associate via $a_{FT}.[F]^2.[T]$ forming a Fst:Tgfb2 complex ($FT$), which dissociates via $-b_{FT}.[FT]$. These terms implement the two-way reaction: $1\,\mathrm{T} + 2\,\mathrm{F} \underset{b_{FT}}{\overset{a_{FT}}{\rightleftharpoons}} 1\,\mathrm{FT}$.

- Fst:Tgfb2 diffuses via $D_{FT}\frac{d^2[FT]}{dx^2}$.

##### Tgfb2:Tgfbr signalling complex:

$$\frac{\partial[RT]}{\partial t} = a_{RT}.[R]^4.[T] - b_{RT}.[RT]$$

Where,

- Active Tgfb2:Tgfbr signalling complexes ($RT$) are formed via $a_{RT}.[R]^4.[T]$. This implements the one-way reaction: $1\,\mathrm{T} + 4\,\mathrm{R} \overset{a_{RT}}{\longrightarrow} 1\,\mathrm{RT}$, whereby one Tgfb2 dimer associates with four Tgfbr monomers (two type I receptors + two type II receptors). This reaction is irreverible since cells internalise and ultimately turn over activated receptor complexes.

- Active Tgfb2:Tgfbr signalling complexes are turned over via $-b_{RT}.[RT]$.

##### R implementation:

The function 'modelA' implements the above system of equations and calculates the rates of change in concentration for each molecular species. The code breaks down into the following steps:

- Fetch the current system state (i.e. molecular concentrations for each 'cell' in the tissue).
- Evaluate the reaction terms for each molecular species.
- Evaluate diffusion terms for diffusible species. Diffusion terms are written for zero-flux boundary conditions. Terms for periodic boundaries are provided but commented out due to considerably longer execution times.
- Combine reaction and diffusion terms to determine rates of change for each species.
- Update a progress bar.
- Return the list of rates.

In [1]:

```
library(deSolve)
library(ReacTran)

## ================
## Model equations
## ================
modelA <- function(time, state, parms) {
  with (as.list(parms), {

    # Generalise this: state[ ( (i-1)*N + 1) : (i * N) ]
    PAX6  <- state[1:N]
    TGFB2 <- state[(N+1):(2*N)]
    FST   <- state[(2*N+1):(3*N)]
    FT    <- state[(3*N+1):(4*N)]
    TGFBR <- state[(4*N+1):(5*N)]
    RT    <- state[(5*N+1):(6*N)]

    ## Reaction terms:
    reacPAX6  <- rep(0, N) # Concentration does not vary throughout simulation
    reacTGFB2 <- (
                  + (aTGFB2 * PAX6 / (PAX6 + 1) ) * (aTGFB2 * PAX6 / (PAX6 + 1) )
                  - (aFT * FST * FST * TGFB2 )
                  + (bFT * FT )
                  - (aRT * TGFBR * TGFBR * TGFBR * TGFBR * TGFB2)
                  - (bTGFB2 * TGFB2)
                 )
    reacFST   <- (
                  + (aFST * (PAX6 / (PAX6 + 1) ) )
                  - (2 * aFT * FST * FST * TGFB2)
                  + (2 * bFT * FT)
                  - (bFST * FST)
                 )
    reacFT    <- (
                  + (aFT * FST * FST * TGFB2)
                  - (bFT * FT)
                 )
    reacTGFBR <- rep(0, N) # Concentration does not vary throughout simulation
    reacRT    <- (
                  + (aRT * TGFBR * TGFBR * TGFBR * TGFBR * TGFB2)
                  - (bRT * RT)
                 )

    ## Diffusion terms - periodic boundary
    ## NOTE: these take longer to compute
    #diffTGFB2 <- tran.1D(C = TGFB2,
    #                     C.up = TGFB2[N],
    #                     C.down = TGFB2[1],
    #                     D = dDimTGFB2,
    #                     dx = xgrid)$dC
    #diffFST   <- tran.1D(C = FST,
    #                     C.up = FST[N],
    #                     C.down = FST[1],
    #                     D = dFST,
    #                     dx = xgrid)$dC
    #diffFT    <- tran.1D(C = FT,
    #                     C.up = FT[N],
    #                     C.down = FT[1],
    #                     D = dFT,
    #                     dx = xgrid)$dC

    ## Diffusion terms - zero-flux boundary
    diffTGFB2 <- tran.1D(C = TGFB2,
                        C.up = TGFB2[1],
                        C.down = TGFB2[N],
                        D = dTGFB2,
                        dx = xgrid)$dC
    diffFST   <- tran.1D(C = FST,
                        C.up = FST[1],
                        C.down = FST[N],
                        D = dFST,
                        dx = xgrid)$dC
    diffFT    <- tran.1D(C = FT,
                        C.up = FT[1],
                        C.down = FT[N],
                        D = dFT,
                        dx = xgrid)$dC

    ## Rates of change = reaction  + diffusion:
    deltaPAX6          = reacPAX6
    deltaTGFB2         = reacTGFB2 + diffTGFB2
    deltaFST           = reacFST   + diffFST
```

```
    deltaFT            = reacFT    + diffFT
    deltaTGFBR         = reacTGFBR
    deltaRT            = reacRT

    setTxtProgressBar(pb, time)

    return (list(c(deltaPAX6, deltaTGFB2, deltaFST, deltaFT, deltaTGFBR, deltaRT)))
  })
}
```

```
Loading required package: rootSolve
Loading required package: shape
```

#### Parameters:

In [2]:

```
## ==================
## Model parameters:
## ==================

## Space:
L      <- 50                              # Length of the 1D tissue
N      <- 100                             # Number of 1D 'cells'
dx     <- L/N                             # Length of one 1D 'cell'
xgridA <- setup.grid.1D(N = N, x.down = L) # Output wanted at these cell positions

## Time:
TIME    <- 1500                   # Duration of the simulation
STEPS   <- 40                     # Number of time-steps
dt      <-  TIME/STEPS            # Size of each time-step
timesA  <- seq(0, TIME, by = dt) # Output wanted at these time intervals

# Setup text progress bar - called by model function
pbA <- txtProgressBar(min = 0, max = TIME, style = 3)

## Rate constants:
parmsA <- c(pb     <- pbA,     # Progress bar
            xgrid  <- xgridA,

            aTGFB2 = 0.39,#0.7,     # Production rate constant for TGFB2 dimer
            bTGFB2 = 0.1,      # Deacy rate constant for TGFB2 dimer
            dTGFB2 = 1,        # Diffusion rate constant for TGFB2 dimer

            aFST   = 1.5,      # Production rate constant for FST
            bFST   = 0.1,      # Decay rate constant for FST
            dFST   = 1,        # Diffusion rate constant for FST

            aFT    = 5,        # Association rate constant for FST:TGFB2 complex
            bFT    = 1,        # Dissociation rate constant for FST:TGFB2 complex
            dFT    = 100,      # Diffusion rate constant for FST:TGBF2 complex

            aRT    = 5,        # Association rate constant for TGFBR:TGFB2 complex
            bRT    = 0.2)      # Decay rate constant for TGFBR:TGFB2 complex
```

```
  |                                                                      |   0%
```

#### Initial Conditions:

The initial conditions describe a situation in which Pax6 expression is restricted to one region within the simulated tissue, corresponding to the distal optic vesicle. This is analagous to the situation in the embryo, where BMPs from the presumptive lens ectoderm promote Pax6 expression in the distal optic vesicle (see main text). Initial distributions of all other molecular species are uniform.

In [3]:

```
## ==================
## Initial conditions:
## ==================
stateA <- c(rep(0.01, N*(1.5/5)), rep(3, N*(2/5)), rep(0.01, N*(1.5/5)),  # PAX6 pre-pattern
            rep(0.1, N),                                                  # TGFB2
            rep(0, N),                                                    # FST
            rep(0, N),                                                    # FT
            rep(1, N),                                                    # TGFBR
            rep(0.0000001, N) )                                           # RT
```

#### Run Simulation:

In [4]:

```
## ==================
# Run the simulation:
## ==================
out <- ode.1D(y         = stateA,
              times     = timesA,
              func      = modelA,
              parms     = parmsA,
              maxsteps  = 1000000,
              nspec     = 6,
              names     = c("PAX6",
                            "TGFB2 Dimer",
                            "FST Monomer",
                            "FST:TGFB2 Heterotetramer",
                            "TGFBR Monomer",
                            "TGFB2:TGFBR Complex"))

# Close the progress bar
close(pb)

# Plot the output
image(out,
      grid       = seq(dx/2, L, dx),
      legend     = TRUE,
      xlab       = "Time",
      ylab       = "1D Space",
      useRaster  = TRUE)


```

```
|=====================================================================  |  99%
```

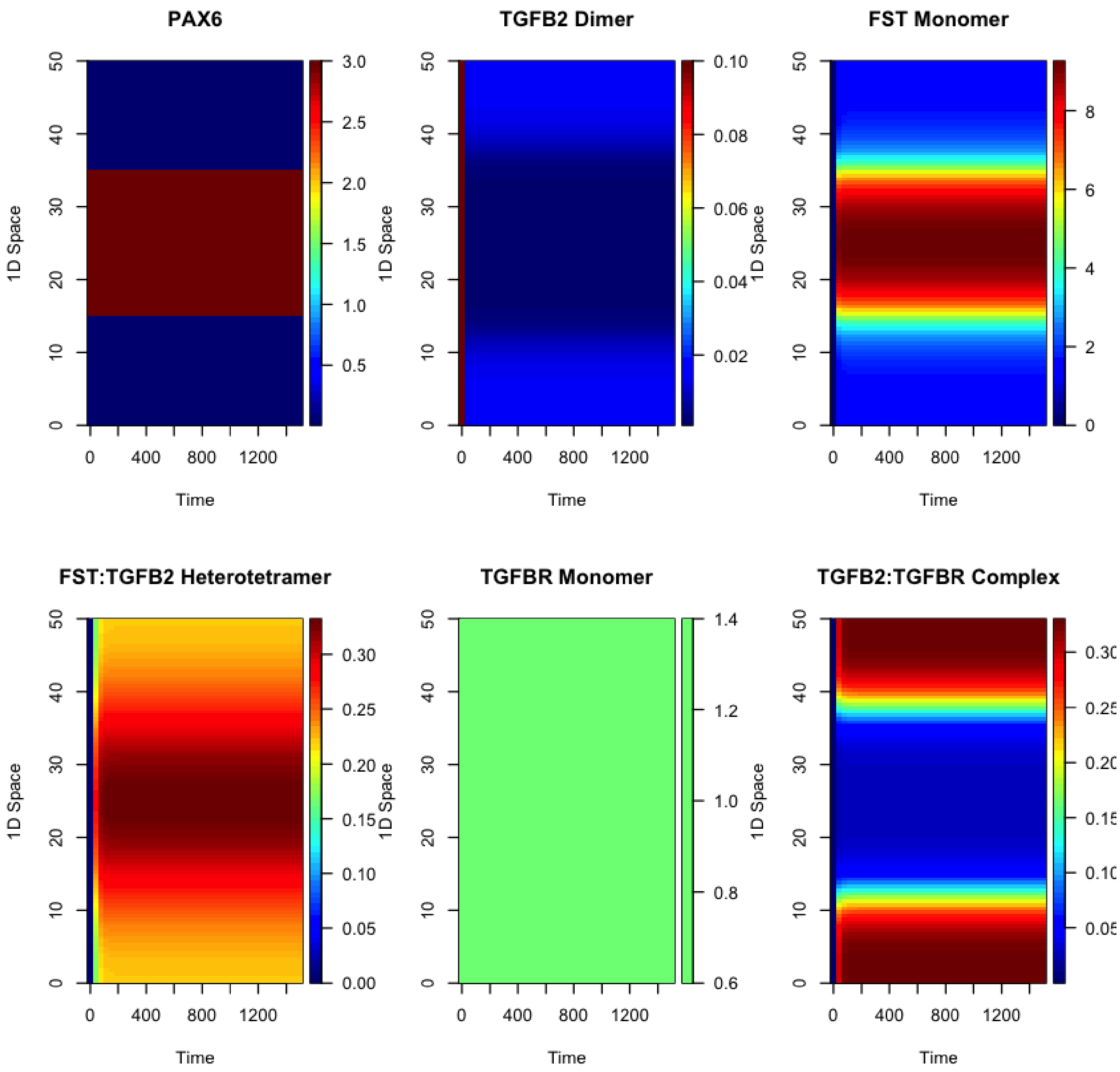

### Reaction-Diffusion Model B:

#### Introduction:

Model B is a direct extension of Model A, whereby Tgfb signalling inhibits Pax6's transcriptional activator function.

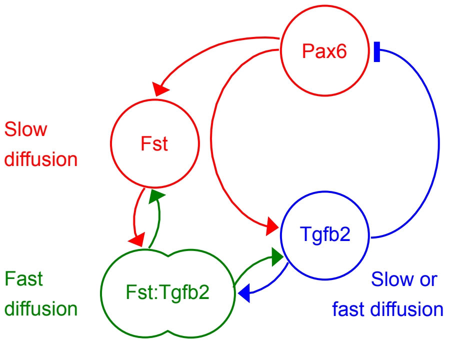


Note: In the following implementation of Model B Pax6 positively autoregulates its own expression. However, this is not strictly necessary for self-organisation to occur. This is because an indirect positive autoregulatory loop already exists; Pax6 acts via Fst to suppress Tgfb2's inhibition of Pax6 within the distal (*Pax6/Fst/Tgfb2*-expressing) pole. This indirect positive autoregulatory loop ("local autocatalysis") combined with lateral activation of Tgfb receptors ("long-range inhibition" of Pax6 function) is sufficient for self-organisation, though some parameter choices may require adjustment form those presented here.

#### Equations:

##### Pax6:

$$\frac{\partial[P]}{\partial t} = a_P \,.\, \frac{[P]}{([P]+1)} \,.\, \frac{1}{[RT]} - b_P \,.\, [P]$$

Where,

- Pax6 ($P$) positively autoregulates via Michaelis-Menton kinetics $\frac{[P]}{[P]+1}$.
- Activated Tgfbr complexes ($RT$) suppress Pax6 function via $\frac{1}{[RT]}$.
- Pax6 is turned over via $-b_P \,.\, [P]$.

##### Tgfb2:

$$\frac{\partial[T]}{\partial t} = \left(a_T \,.\, \frac{[P]}{([P]+1)} \,.\, \frac{1}{[RT]}\right)^2 - a_{FT} \,.\, [F]^2 \,.\, [T] + b_{FT} \,.\, [FT] - a_{RT} \,.\, [R]^4 \,.\, [T] - b_T \,.\, [T] + D_T \frac{d^2[T]}{dx^2}$$

Where,

- Pax6 ($P$) positively regulates Tgfb2 expression via Michaelis-Menton kinetics $\frac{[P]}{[P]+1}$.

- Activated Tgfbr complexes ($RT$) suppress Pax6 function via $\frac{1}{[RT]}$.

- Tgfb2 is produced as a dimer, hence $\left(a_T \,.\, \frac{[P]}{([P]+1)} \,.\, \frac{1}{[RT]}\right)^2$.

- Tgfb2 is sequestered in the formation of Fst:Tgfb2 complexes ($FT$) via $-a_{FT} \,.\, [F]^2 \,.\, [T]$, and released again via $+b_{FT} \,.\, [FT]$. This implements the two-way reaction: $1\,\mathrm{T} + 2\,\mathrm{F} \underset{b_{FT}}{\overset{a_{FT}}{\rightleftharpoons}} 1\,\mathrm{FT}$, whereby one Tgfb2 dimer reversibly associates with two Fst monomers.

- Tgfb2 is irreversibly consumed in the formation of activated receptor complexes ($RT$) via $-a_{RT} \,.\, [R]^4 \,.\, [T]$. This implements the one-way reaction: $1\,\mathrm{T} + 4\,\mathrm{R} \xrightarrow{a_{RT}} 1\,\mathrm{RT}$, whereby one Tgfb2 dimer associates with four Tgfbr monomers (two type I receptors + two type II receptors). This reaction is irreverible since cells internalise and ultimately turn over activated receptor complexes.

- Tgfb2 is turned over via $-b_T \,.\, [T]$.

- Tgfb2 diffuses via $D_T \frac{d^2[T]}{dx^2}$.

##### Fst:

$$\frac{\partial[F]}{\partial t} = a_F \,.\, \frac{[P]}{([P]+1)} \,.\, \frac{1}{[RT]} - 2.a_{FT} \,.\, [F]^2 \,.\, [T] + 2.b_{FT} \,.\, [FT] - b_F \,.\, [F] + D_F \frac{d^2[F]}{dx^2}$$

Where,

- Pax6 ($P$) positively regulates Fst expression via Michaelis-Menton kinetics $\frac{[P]}{[P]+1}$.

- Activated Tgfbr complexes ($RT$) suppress Pax6 function via $\frac{1}{[RT]}$.

- Pairs of Fst monomers are consumed in the formation of Fst:Tgfb2 complexes ($FT$) via $-2.a_{FT}.[F]^2.[T]$, and released again via $+2.b_{FT}.[FT]$. This implements the two-way reaction: $1\,\mathrm{T} + 2\,\mathrm{F} \underset{b_{FT}}{\overset{a_{FT}}{\rightleftharpoons}} 1\,\mathrm{FT}$, whereby one Tgfb2 dimer reversibly associates with two Fst monomers.

- Fst is turned over via $-b_F.[F]$.

- Fst diffuses via $D_F \frac{d^2[F]}{dx^2}$.

#### Fst:Tgfb2 complex:

$$\frac{\partial[FT]}{\partial t} = a_{FT}.[F]^2.[T] - b_{FT}.[FT] + D_{FT}\frac{d^2[FT]}{dx^2}$$

Where,

- A pair of Fst monomers and a single Tgfb2 dimer associate via $a_{FT}.[F]^2.[T]$ forming a Fst:Tgfb2 complex, which dissociates via $-b_{FT}.[FT]$. These terms implement the two-way reaction: $1\,\mathrm{T} + 2\,\mathrm{F} \underset{b_{FT}}{\overset{a_{FT}}{\rightleftharpoons}} 1\,\mathrm{FT}$.

- Fst:Tgfb2 diffuses via $D_{FT}\frac{d^2[FT]}{dx^2}$.

#### Activated Tgfbr complex:

$$\frac{\partial[RT]}{\partial t} = a_{RT}.[R]^4.[T] - b_{RT}.[RT]$$

#### R implementation:

The function 'modelB' implements the above system of equations and calculates the rates of change in concentration for each molecular species. The code breaks down into the following steps:

- Fetch the current system state (i.e. molecular concentrations for each 'cell' in the tissue).
- Evaluate the reaction terms for each molecular species.
- Evaluate diffusion terms for diffusible species. Diffusion terms are written for zero-flux boundary conditions. Terms for periodic boundaries are provided but commented out due to considerably longer execution times.
- Combine reaction and diffusion terms to determine rates of change for each species.
- Update a progress bar.
- Return the list of rates.

In [5]:

```r
library(deSolve)
library(ReacTran)

## ================
## Model equations
## ================
modelB <- function(time, state, parms) {
  with (as.list(parms), {
    # Generalise this: state[ ( (i-1)*N + 1) : (i * N) ]
    PAX6  <- state[1:N]
    TGFB2 <- state[(N+1):(2*N)]
    FST   <- state[(2*N+1):(3*N)]
    FT    <- state[(3*N+1):(4*N)]
    TGFBR <- state[(4*N+1):(5*N)]
    RT    <- state[(5*N+1):(6*N)]

    ## Reaction terms:
    reacPAX6   <- (
                 + (aPAX6 * PAX6) / (RT * (PAX6 + 1) )
                 - (bPAX6 * PAX6)
                 )
    reacTGFB2 <- (
                 + (aTGFB2 * PAX6 / (RT * (PAX6 + 1) ) ) * (aTGFB2 * PAX6 / (RT * (PAX6 + 1) ) )
                 - (aFT * FST * FST * TGFB2 )
                 + (bFT * FT)
                 - (aRT * TGFBR * TGFBR * TGFBR * TGFBR * TGFB2)
                 - (bTGFB2 * TGFB2)
                 )
    reacFST   <- (
                 + (aFST * PAX6) / (RT * (PAX6 + 1) )
                 - (2 * aFT * FST * FST * TGFB2)
                 + (2 * bFT * FT)
                 - (bFST * FST)
                 )
    reacFT    <- (
                 + (aFT * FST * FST * TGFB2)
                 - (bFT * FT)
                 )
    reacTGFBR <- rep(0, N) # Concentration does not vary throughout simulation
    reacRT    <- (
                 + (aRT * TGFBR * TGFBR * TGFBR * TGFBR * TGFB2)
                 - (bRT * RT)
                 )

    ## Diffusion terms - periodic boundary
    ## NOTE: these take longer to compute
    #diffTGFB2 <- tran.1D(C = TGFB2,
    #                      C.up = TGFB2[N],
    #                      C.down = TGFB2[1],
    #                      D = dDimTGFB2,
    #                      dx = xgrid)$dC
    #diffFST   <- tran.1D(C = FST,
    #                      C.up = FST[N],
    #                      C.down = FST[1],
    #                      D = dFST,
    #                      dx = xgrid)$dC
    #diffFT    <- tran.1D(C = FT,
    #                      C.up = FT[N],
    #                      C.down = FT[1],
    #                      D = dFT,
    #                      dx = xgrid)$dC

    ## Diffusion terms - zero-flux boundary
    diffTGFB2 <- tran.1D(C = TGFB2,
                        C.up = TGFB2[1],
                        C.down = TGFB2[N],
                        D = dTGFB2,
                        dx = xgridB)$dC
    diffFST   <- tran.1D(C = FST,
                        C.up = FST[1],
                        C.down = FST[N],
                        D = dFST,
                        dx = xgridB)$dC
    diffFT    <- tran.1D(C = FT,
                        C.up = FT[1],
                        C.down = FT[N],
                        D = dFT,
                        dx = xgridB)$dC

    ## Rates of change = Reaction  + Diffusion:
    deltaPAX6          = reacPAX6
```

```
    deltaTGFB2          = reacTGFB2 + diffTGFB2
    deltaFST            = reacFST   + diffFST
    deltaFT             = reacFT    + diffFT
    deltaTGFBR          = reacTGFBR
    deltaRT             = reacRT

    setTxtProgressBar(pb, time)

    return (list(c(deltaPAX6, deltaTGFB2, deltaFST, deltaFT, deltaTGFBR, deltaRT)))
  })
}
```

#### Parameters:

In [6]:

```
## ==================
## Model parameters:
## ==================

## Space:
L      <- 100                                # Length of the 1D tissue
N      <- 100                               # Number of 1D 'cells'
dx     <- L/N                               # Length of one 1D 'cell'
xgridB <- setup.grid.1D(N = N, x.down = L) # Output wanted at these cell positions

## Time:
TIME    <- 10000                            # Duration of the simulation
STEPS   <- 40                               # Number of time-steps
dt      <-  TIME/STEPS                      # Size of each time-step
timesB  <- seq(0, TIME, by = dt)            # Output wanted at these time intervals

# Setup text progress bar - called by model function
pbB <- txtProgressBar(min = 0, max = TIME, style = 3)

## Rate constants:
parmsB <- c(pb     <- pbB,    # Progress bar
            xgrid  <- xgridB,

            aPAX6  = 0.3,     # Production rate constant for PAX6
            bPAX6  = 0.1,     # Decay rate constant for PAX6

            aTGFB2 = 0.39,    # Production rate constant for TGFB2 dimer
            bTGFB2 = 0.1,     # Deacy rate constant for TGFB2 dimer
            dTGFB2 = 1,       # Diffusion rate constant for TGFB2 dimer

            aFST   = 1.5,     # Production rate constant for FST
            bFST   = 0.1,     # Decay rate constant for FST
            dFST   = 1,       # Diffusion rate constant for FST

            aFT    = 5,       # Association rate constant for FST:TGFB2 complex
            bFT    = 1,       # Dissociation rate constant for FST:TGFB2 complex
            dFT    = 100,     # Diffusion rate constant for FST:TGBF2 complex

            aRT    = 5,       # Association rate constant for TGFBR:TGFB2 complex
            bRT    = 0.2)     # Decay rate constant for TGFBR:TGFB2 complex
```

```
  |                                                                      |   0%
```

#### Initial Conditions:

The initial conditions describe a situation in which Pax6 expression is expressed at a low level throughout the entire tissue. This inital distribution is assumed to be noisey with small fluctuations around the average expression level from one cell to the next.

This is analagous to a situation in which there is an absence of well ordered positional information from interactions with neighbouring eye tissues (e.g. presumptive lens ectoderm, peri-ocular mesenchyme etc.) such as in retinal organoid cultures. Initial distributions of all other molecular species are uniform.

In [7]:

```
## ==================
## Initial conditions:
## ==================
stateB <- c(jitter( rep( (1.9)/2 , N), (0.1)/2, (0.1)/2), #PAX6 - no pre-pattern
            rep(0.1, N),                                  # TGFB2
            rep(0, N),                                    # FST
            rep(0, N),                                    # FT
            rep(1, N),                                    # TGFBR
            rep(0.0000001, N) )                           # RT
```

**Run Simulation:**

In [8]:

```
## =================
# Run the simulation:
## =================
out <- ode.1D(y         = stateB,
              times     = timesB,
              func      = modelB,
              parms     = parmsB,
              maxsteps  = 1000000,
              nspec     = 6,
              names     = c("PAX6",
                            "TGFB2 Dimer",
                            "FST Monomer",
                            "FST:TGFB2 Heterotetramer",
                            "TGFBR Monomer",
                            "TGFB2:TGFBR Complex"))

# Close the progress bar
close(pbB)

# Plot the output
image(out,
      grid       = seq(dx/2, L, dx),
      legend     = TRUE,
      xlab       = "Time",
      ylab       = "1D Space",
      useRaster  = TRUE)


```

```
|======================================================================| 100%
```

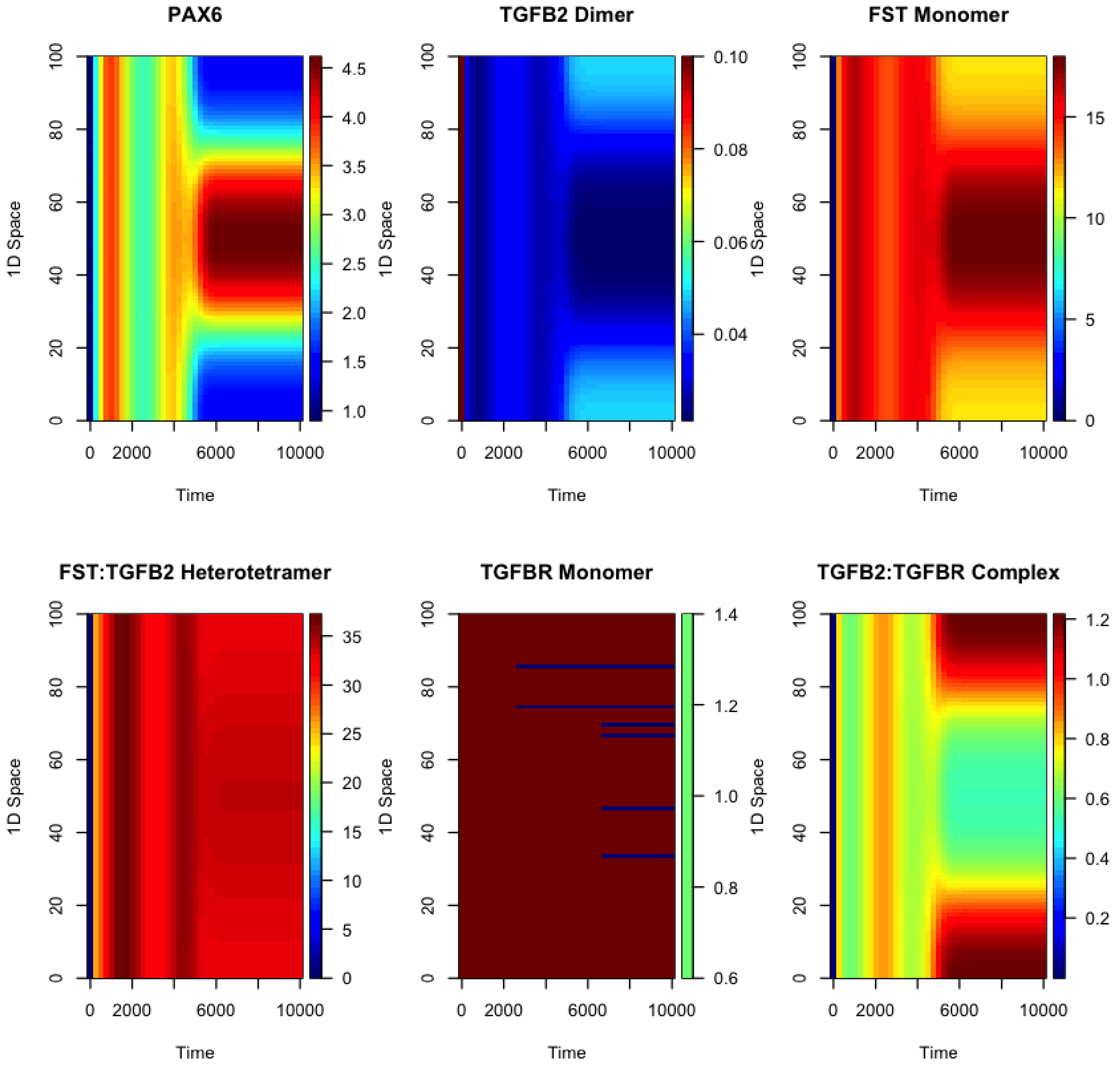
